# Structural Interaction Fingerprints and Machine Learning for predicting and explaining binding of small molecule ligands to RNA

**DOI:** 10.1101/2023.01.11.523582

**Authors:** Natalia A. Szulc, Zuzanna Mackiewicz, Janusz M. Bujnicki, Filip Stefaniak

**Affiliations:** Laboratory of Bioinformatics and Protein Engineering, International Institute of Molecular and Cell Biology in Warsaw, 4 Ks. Trojdena Str., 02-109 Warsaw, Poland; Laboratory of Protein Metabolism, International Institute of Molecular and Cell Biology in Warsaw, 4 Ks. Trojdena Str., 02-109 Warsaw, Poland; Laboratory of RNA Biology - ERA Chairs Group, International Institute of Molecular and Cell Biology in Warsaw, 4 Ks. Trojdena Str., 02-109 Warsaw, Poland

**Keywords:** RNA, small molecules, Structural Interaction Fingerprint, machine learning, explainable artificial intelligence, XAI

## Abstract

Ribonucleic acids (RNA) play crucial roles in living organisms as they are involved in key processes necessary for proper cell functioning. Some RNA molecules, such as bacterial ribosomes and precursor messenger RNA, are targets of small molecule drugs, while others, e.g., bacterial riboswitches or viral RNA motifs are considered as potential therapeutic targets. Thus, the continuous discovery of new functional RNA increases the demand for developing compounds targeting them and for methods for analyzing RNA—small molecule interactions. We recently developed fingeRNAt - a software for detecting non-covalent bonds formed within complexes of nucleic acids with different types of ligands. The program detects several non-covalent interactions, such as hydrogen and halogen bonds, ionic, Pi, inorganic ion-and water-mediated, lipophilic interactions, and encodes them as computational-friendly Structural Interaction Fingerprint (SIFt). Here we present the application of SIFts accompanied by machine learning methods for binding prediction of small molecules to RNA targets. We show that SIFt-based models outperform the classic, general-purpose scoring functions in virtual screening. We discuss the aid offered by Explainable Artificial Intelligence in the analysis of the binding prediction models, elucidating the decision-making process, and deciphering molecular recognition processes.

**Key Points:**

- Structural Interaction fingerprints (SIFts), combined with machine learning, were successfully used to develop activity models for ligands binding to RNA.
- SIFt-based models outperformed the classic, general-purpose scoring functions in virtual screening.
- Explainable Artificial Intelligence allowed us to understand the decision-making process and decipher molecular recognition processes in the analysis of RNA—ligand binding activity models.
- We provide a benchmark dataset based on ligands with known or putative binding activity toward six RNA targets. It can be readily used by the scientific community to test new algorithms of virtual screening on RNA—ligand complexes.

## INTRODUCTION

Ribonucleic acids (RNA) are essential bioorganic molecules present in every living organism. Apart from messenger RNA (mRNA), many non-coding RNA (ncRNA) also play a crucial role in the cell, as they build large macromolecular machines, deliver amino acids to ribosomes, or regulate different molecular processes, e.g., by acting as binding sites. While the majority of drugs target proteins [1], only about 1.5% of the human genome consists of protein-coding sequences [2]. On the other hand, there is growing evidence of the functional role of ncRNA and their involvement in a plethora of diseases [3–5]. Also, many nucleic acids, or at least segments/domains within their sequence, possess the ability to adopt tertiary structures and have grooves capable of binding small molecules, making them attractive targets for small molecule drugs [6,7].

As experimental approaches provide the most valuable data of biological activity for small molecule ligands, their high-throughput usage is restricted due to the tremendous cost, as well as the time- and labor-consuming nature of the process. For this reason, the computational virtual screening technique is widely used, as it allows to score and compare thousands of potentially binding ligands much faster and cheaper than can be done experimentally. Thus, effective scoring, i.e., predicting their activity (in terms of binding, as biological activity can only be validated experimentally), is the ultimate challenge of rational drug discovery, as it allows for narrowing the number of compounds for further assays. Molecular docking is the classical method used in numerous studies to predict the favored geometric orientation of the receptor-ligand complex and to virtually screen a database of thousands of small molecule compounds to select those with the highest predicted binding activity towards the receptor for further optimization [8,9].

Numerous computer programs exist for molecular docking. They share common steps of ligand sampling and scoring and have similar drawbacks arising from the fact that more nature-like approximations of receptor-ligand flexible conformation entail a high number of degrees of freedom, thus requiring exhaustive computations. Also, scoring functions used to assess molecular docking results have limited accuracy and may not correctly recognize binding ligands. Recently developed strategies based on machine learning (ML) may alleviate these obstacles and provide new frameworks for predicting active ligands [10,11]. Also, the recent advance in the field of Explainable Artificial Intelligence (XAI) may facilitate the understanding of decisions made by predictive ML models and support decisions made by medicinal chemists [12,13].

Molecular fingerprints translate two- or three-dimensional structural information into one-dimensional vectors. These are machine-readable entities suitable for similarity calculation, modeling, data mining, or ML (for a recent review on molecular fingerprints, see [14,15], and for their applications in ML, see [16]). Structural Interaction Fingerprint (SIFt) is a molecular fingerprint type first described by Deng *et al*. for analyzing protein-ligand complexes [17]. It is a binary string representing the presence of specific molecular interactions between the receptor’s residues and the ligand. SIFts have been used in studying protein-ligand complexes and were successfully applied in a small molecule drug discovery: selectivity profiling, new target prediction, elucidating the binding mechanisms, or developing new scoring functions [18–20]. SIFts enable filtering virtual chemical libraries to select only compounds with similar interaction interfaces to molecules with experimentally validated biological activities [17,21,22]. We recently showed that SIFts can also be successfully applied for the analysis of RNA—ligand complexes: comparing the binding mode in experimentally solved complexes, clustering of docking poses, and selecting ligands forming a similar interaction network with the RNA receptor [23].

However, since most virtual screening methods focus on proteins, there is a scarcity of nucleic acids-oriented approaches. The goal of this work was to determine if SIFts combined with ML techniques can be applied to predict the binding activity of small molecules to RNA targets. As there is no widely-available and up-to-date database of compounds with known binding activity toward RNA targets, for the purpose of testing virtual screening procedures, we composed a benchmark set based on the literature search, using the methodology previously described for protein and RNA targets [23,24], freely available at https://github.com/filipsPL/fingernat-ml [25]. To calculate SIFts, we used our recently published open-source nucleic acid-oriented software fingeRNAt [23]. In our analysis, we compared the performance of the SIFt-based method with the state-of-the-art, RNA-oriented scoring functions. Finally, using the example of human immunodeficiency virus type 1 (HIV-1) trans-activation response element (TAR) RNA, we applied XAI methods to elucidate the structural factors determining the ligand binding.

Together, our work is the first described protocol for predicting ligand activity towards RNA receptors with molecular docking and explainable SIFt- and ML-based scoring techniques.

## RESULTS AND DISCUSSION

To test and compare the predictive power of various ML methods for RNA—small molecule binding, six RNA targets were selected: (i) HIV-1 TAR, (ii) thiamine pyrophosphate (TPP) riboswitch, (iii) flavin mononucleotide (FMN) riboswitch, (iv) adenine riboswitch, (v) guanine riboswitch, and (vi) guanine riboswitch with the C74U mutation (guanine riboswitch with C74U mutation was distinguished as a separate target because of differences in ligand specificity resulting from this nucleotide change [26]). Information on chemical structures and binding properties of small molecule ligands were taken from the literature and experimentally solved structures deposited in the RCSB Protein Data Bank (PDB). To avoid discrepancies resulting from different experimental techniques used to determine ligands binding, we used a binary class to describe their affinity toward RNA target: *binders*, for molecules binding relatively strongly, and *non-binders*, for those with weak or non-detectable binding. For each RNA target, a structurally diversified set of ligands was prepared. As the information on non-binding ligands is sparse in the literature reports, putative non-binding ligands were generated using a well-established DUD-E methodology ([24]; for the procedure details and a number of ligands in the datasets, see Materials and Methods).

For all of the investigated targets, multiple experimentally determined structures are deposited in the RCSB PDB. Therefore, for each RNA target, three distinct structures (but with an identical sequence of the binding pocket within 8 Å from the experimental ligand) were selected for further analysis (see Table 1 from Materials and Methods). Next, the ligands were docked to their targets using the rDock docking software [27]. SIFts were calculated for three top-scored poses for each complex of ligand and its respective RNA target (see Materials and Methods and Fig 5).

**Table 1.** Molecular RNA targets used for benchmarking, PDB IDs of experimentally solved structures used for docking and SIFts calculation, the number of ligands collected from the literature, and the final composition of ligand sets after adding decoys.

| RNA target | PDB IDs | Number of collected ligands | Final dataset (with DUD-E generated decoys) |  |
| --- | --- | --- | --- | --- |
|  |  |  | *Binders* | *Non-binders* |
| Adenine riboswitch | 1Y26, 4TZX, 4XNR | 18 | 7 | 361 |
| FMN riboswitch | 2YIE, 5KX9, 6BFB | 30 | 15 | 750 |
| Guanine riboswitch | 3GAO, 3GER, 4FE5 | 79 | 30 | 1510 |
| Guanine riboswitch with C74U mutation | 2XNW, 2XNZ, 2XO1 | 25 | 14 | 714 |
| HIV-1 TAR | 1UTS, 1UUI, 6MCE | 139 | 30 | 1478 |
| TPP riboswitch | 2GDI, 4NYA, 4NYD | 73 | 30 | 1510 |
| **Total** |  | **364** | **126** | **6323** |

### CatBoost, Extra Trees, and LightGBM are the most effective ML methods in predicting ligand binding

First, we checked the virtual screening performance of the ML algorithms, whose inputs were SIFts for each RNA—ligand complex with assigned ligand activity toward a given RNA target as a binary class (*binder/non-binder*). Six ML classifiers were evaluated at their default setup, with the optional feature selection preprocessing step and/or stacking (see Materials and Methods).

All tested ML methods performed better than the Baseline predictor (the negative control - which returns the most frequent label from the training data), with the average AUROC (area under the receiver operating characteristic curve, calculated from 3-fold cross validation results) ranging from 0.81 (Decision Tree) to 0.97 (LightGBM), and median AUROC values ranging from 0.79 (Decision Tree) to 0.99 (CatBoost; for the performance summary of all examined methods, see Fig 1). The performance varies from one RNA target to another, with the lowest average AUROC value for HIV-1 TAR (0.83) and the highest for TPP riboswitch (0.95). This indicates that binding predictions for some molecular targets are more challenging with the presented method. It may result from an inability of the docking program to correctly predict the binding mode of the ligands, the fact that small molecules bind to the distinct binding pocket of the given RNA, or that the RNA undergoes a conformational change during the ligand binding. However, even for HIV-1 TAR RNA, for which we observed the worst average performance, ML methods with satisfactory AUROC values can still be selected (LightGBM with SelectedFeatures and RandomForest with SelectedFeatures and Stacked, with AUROC values equal to 0.92 and 0.92, respectively).

**Figure 1.**
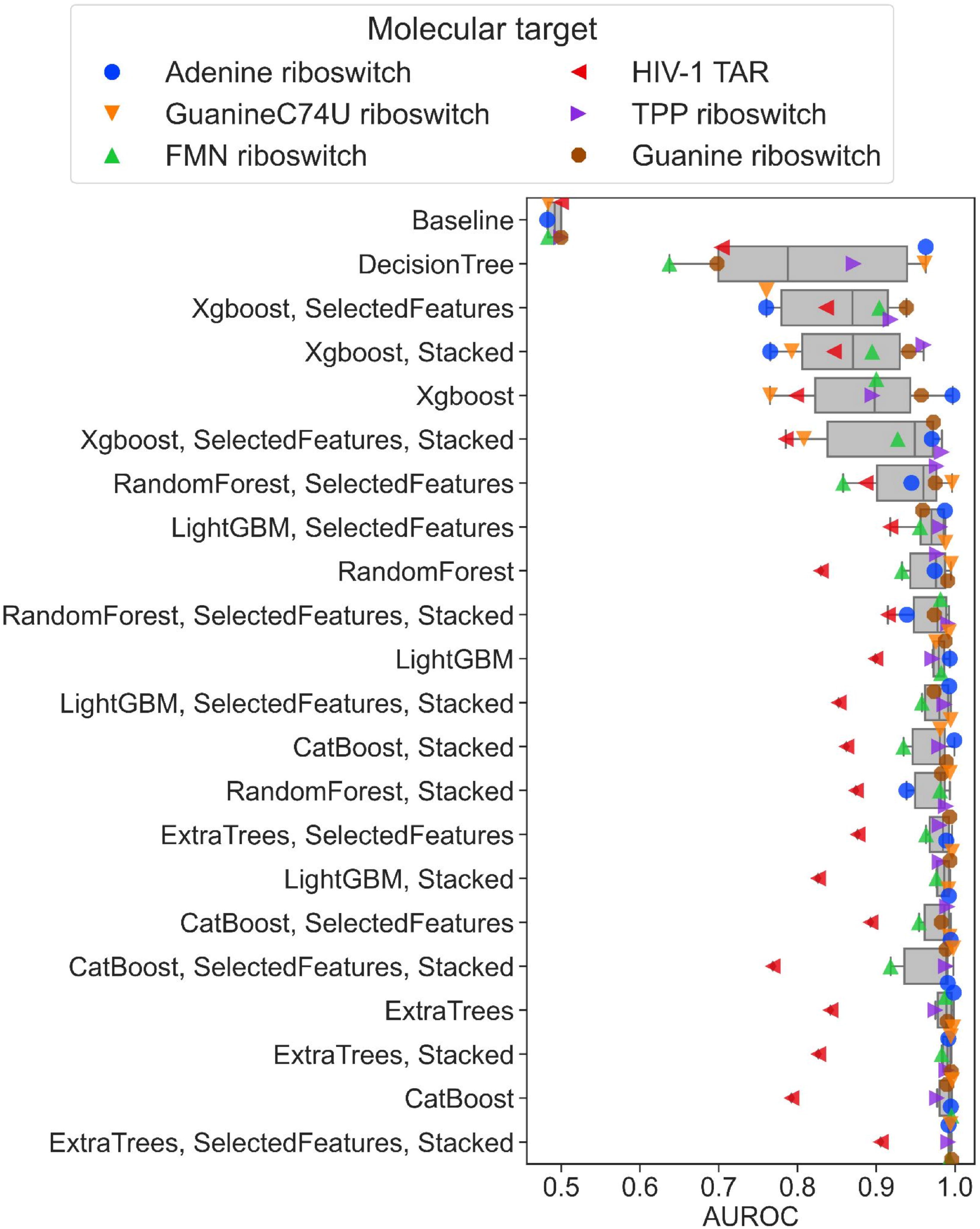
The performance of the ML methods. The performance is expressed as AUROC values (calculated from the 3-fold cross-validation results) and shown as a box-and-whisker plot. Individual AUROC values for each molecular target are overlaid as color- and shape-coded points.

We observed that - on average - the additional pre- and post-processing steps slightly decreased the performance of the ML methods when applied individually (with the average AUROC values for all methods and targets equal to 0.94 and 0.94 for feature selection and stacking, respectively, compared to 0.95 for methods without pre- and post-processing) but the exact influence depends on the method. Feature Selection improved two methods (CatBoost and Extra Trees) and worsened three methods (LightGBM, Random Forest, and XGBoost), while stacking improved one method (Random Forest), worsened three methods (Extra Trees, LightGBM, and XGBoost), and did not influence one method (CatBoost) (Fig 1). Nevertheless, all differences in observed AUROC values were not spectacular. Feature selection and stacking, when combined, also had a minor influence on the observed AUROC values. On average, it slightly increased the average performance (AUROC value 0.95). The accuracy was better for three methods (Extra Trees, Random Forest, and XGBoost) and worsened for two methods (CatBoost, LightGBM), but the differences in AUROC values were still minor.

### SIFts, in combination with ML methods, outperform classic scoring functions in ligand binding predictions

As shown above, SIFts combined with ML effectively predict ligand binding activity toward RNA targets. We compared these methods with the performance of classical general-purpose scoring functions. For that, we compared the screening metrics (AUROC, Boltzmann-enhanced discrimination of receiver operating characteristic - BEDROC, and Enrichment Factor for the top-scored 10% of ligands - EF_10%_; [28]) for the three best ML methods trained on SIFts (namely CatBoost, Extra Trees, and LightGBM), three classic RNA—ligand scoring functions (AnnapuRNA [29], LigandRNA [30], and rDock [31]) and one protein-ligand scoring function (rf-score-vs, as a negative control [32]). The results are summarized in Fig 2.

**Figure 2.**
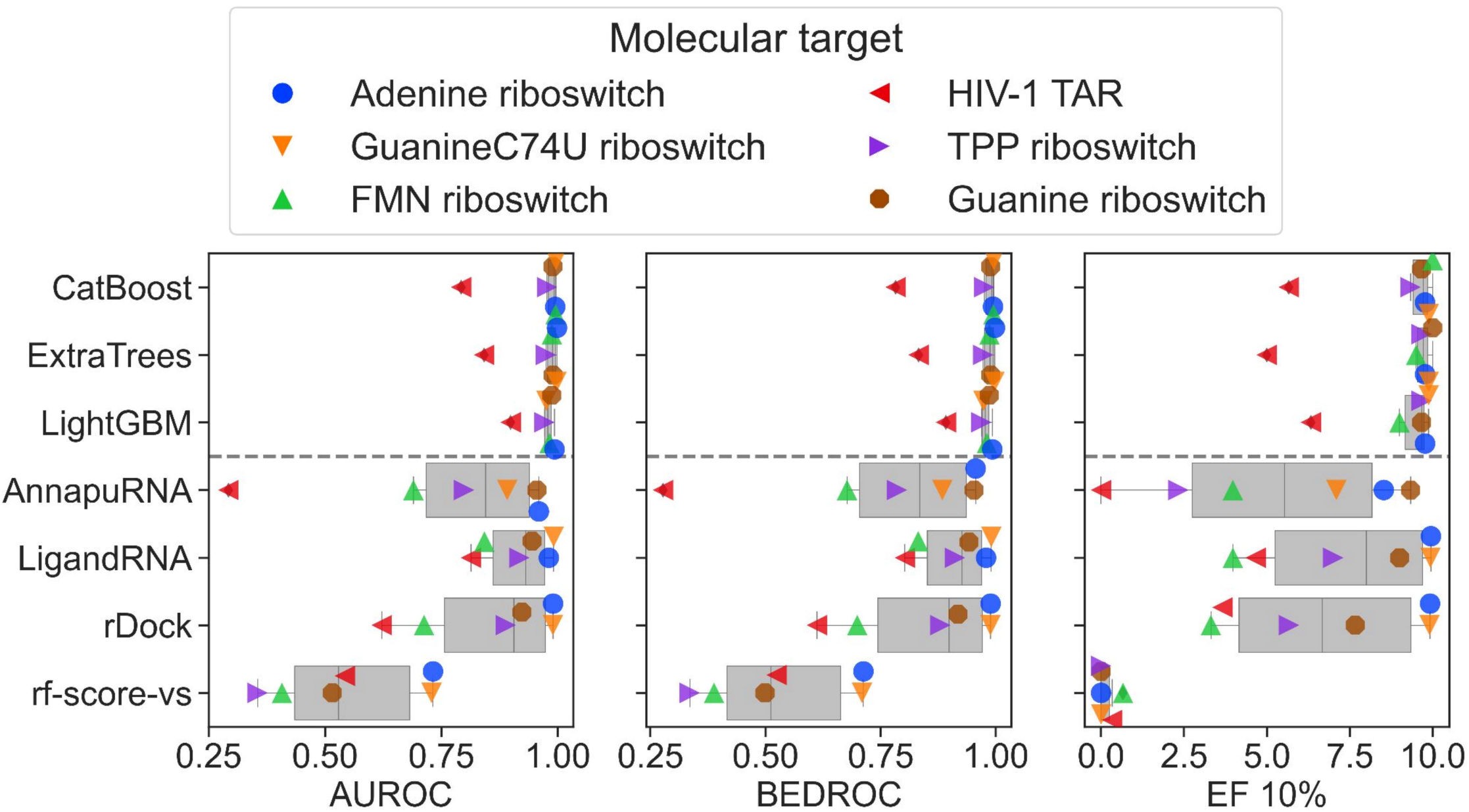
Comparison of AUROC values between ML methods. The performance of the three best ML methods (CatBoost, Extra Trees, and LightGBM, with no feature selection nor stacking applied, calculated from the 3-fold cross-validation results) trained on the SIFts (upper part of the plots) is compared with the performance of the scoring functions (AnnapuRNA, LigandRNA, rDock, and rf-score-vs; the bottom part of the plots). The distribution of AUROC, BEDROC, and EF_10%_ values are represented on box-and-whisker plots, and the individual performance for six molecular targets is overlaid as color- and shape-coded points. The performance of scoring functions is displayed for the best performance among the three structures for each RNA.

The data show that the average performance of SIFts-based scoring functions is superior to the examined classic scoring functions, although the results are highly dependent on the molecular target. For example, for the adenine riboswitch, the SIFts methods reached AUROC values ranging from 0.99 to 1.00, while classic RNA—ligand scoring functions have only slightly worse performance (ranging from 0.96 to 0.99). This almost-ideal performance may be partially caused by the fact that the chemical space of binding ligands of adenine and guanine riboswitches is limited to heterocyclic compounds of similar size, which are also structurally similar to the experimentally determined ligands. This increases the probability of finding the native or near-native conformation of the complex by the docking program and results in satisfactory screening performance for both our method and classic RNA—ligand scoring functions. It should be noted that our datasets for guanine and adenine riboswitches also contain heterocyclic experimentally-confirmed non-binding molecules; thus, the *binder*/*non-binder* recognition process goes beyond simple structural rules.

Similarly, the performance of the SIFts methods for FMN riboswitch is also very good (with AUROC ranging from 0.98 to 1.00), while for the scoring functions, it is much worse (AUROC ranging from 0.69 to 0.84), and for the protein dedicated rf-score-vs AUROC reaches value 0.41 (which means worse than the random selection of the binders). For a single target - HIV-1 TAR RNA, we observed the decreased performance of all examined scoring functions (for SIFts methods, AUROC values range from 0.79 to 0.90, for classical functions from 0.29 to 0.81, for protein scoring function equals 0.54). This shows that the molecular recognition process for such “difficult” cases may be beyond the relatively simple interaction model utilized by the current scoring functions (both described in this work and developed earlier) and cannot recapitulate the complicated binding phenomena observed in real life.

As expected, for all the cases, the performance of the protein-ligand scoring function is much worse than for ML based methods or classical scoring functions, with the AUROC value ranging from 0.36 for TPP riboswitch to 0.73 for adenine riboswitch.

The obtained benchmark results indicate that the SIFts-based “local” scoring functions, which are trained on the binding activity data for a given molecular target, perform visibly better than the general-purpose scoring functions; however, this difference is not statistically significant. Also, our data pinpoint the need for further development and tests of the scoring functions dedicated to the RNA—ligands complexes, as the performance of the protein scoring functions may be very poor, frequently offering a random or worse-than-random selection of the ligands. The limitation is, however, the availability of the training data required to build the interaction model.

### Explainable ML models

Despite the great accuracy and support offered by many ML models in drug discovery and molecular biology, their application is limited by the lack of transparency on how and why the decisions are made [33]. To overcome this little confidence in Artificial Intelligence systems, XAI became crucial, especially for application in biomedical, medical, and healthcare studies. Also, in drug design applications, interpretable and explainable ML models offer support in understanding the binding phenomena, interpreting the landscape of structure-activity relationships in the inspected pool of compounds, or deciphering complex drug-drug interactions [34–36].

We investigated if the interpretation of binding prediction of the ML models built on SIFts can give insights into the factors important for the small molecule ligand binding to the RNA. Similar work has been recently presented for SIFts and a protein target - dopamine D4 receptor - and was proven useful in indicating important factors for ligand binding [37]. In our study, we used a locally interpretable explanatory method termed SHapley Additive exPlanations (SHAP), derived from cooperative game theory and applicable to multiple ML problems in different fields [38]. For each prediction (each ligand) and each feature used in the predictive model, a SHAP value is assigned to quantify how this particular feature influences the prediction (in this work: the probability that the ligand belongs to the *binders* class). For this experiment, we built an ML model using one of the best-performing methods - CatBoost, for HIV-1 TAR class-balanced data. This RNA was chosen to demonstrate the usability of the proposed approach for more difficult but medically-relevant molecular target. For clarity of data presentation, we used SIFts with the basic interactions subset (hydrogen and halogen bonds, cation-anion, Pi-cation, Pi-anion, Pi-stacking, and lipophilic interactions; see [23]), derived for the top-scored pose of each ligand docked to a single RNA structure (PDB ID: 1UTS, chosen arbitrarily). In addition to the SHAP method, we estimated the variable importances for the generated ML model using the CatBoost classifiers: CatBoost Prediction Values Change (indicating how much, on average, the prediction changes if the feature value changes), and the CatBoost Loss Function Change (indicating the difference between the loss value of the model with this feature and without it). The feature importance summary is presented in Fig 3.

**Figure 3.**
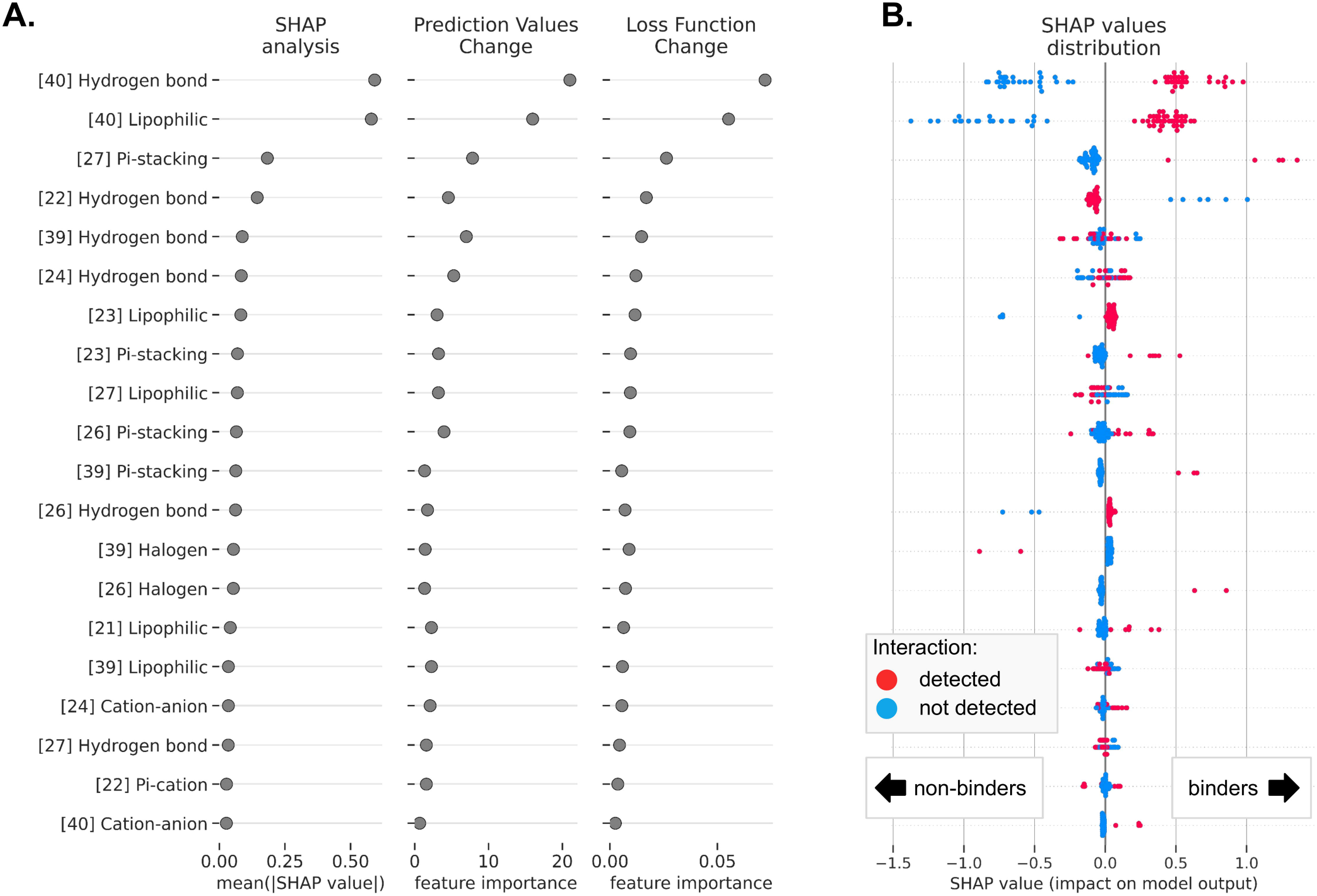
Feature importance analysis for the HIV-1 TAR activity binding model trained with the CatBoost classifier on SIFts. Each row indicates a selected interaction type formed with a given residue (number in square brackets). **(A)** The 20 top-ranked interactions and their importance for the model performance tested with the SHAP method (the contribution of each feature to the prediction of the model - mean absolute SHAP value), the CatBoost Prediction Values Change (indicating how much, on average, the prediction changes if the feature value changes), and the CatBoost Loss Function Change (indicating the difference between the loss value of the model with this feature and without it). **(B)** The distribution of SHAP values for all ligands in the data set, with color-coded presence/absence of the interactions of the given type (red - the interaction was detected, blue - the interaction was not detected). SHAP values show how forming the given interaction increases or decreases the likelihood of a compound being classified as a *binder*.

According to all three methods, the most important for the binding activity model are interactions with residue 40 (hydrogen bonds and lipophilic interactions) and residue 27 (Pi-stacking interaction) (Fig 3A). The forming of these interactions positively contributes to the model output and increases the predicted probability of binding activity for ligands (Fig 3B). This is in agreement with the experimental data deposited in the literature [39]. The subsequent, less important interactions with a positive contribution to the model output are non-covalent bonds with residues 23 (lipophilic and Pi-stacking; this residue is essential for binding of native substrate, Tat protein, to HIV-1 TAR RNA, and is highly conserved among different subtypes of HIV-1 and HIV-2, and simian immunodeficiency virus (SIV) [40], thus utilizing it for ligand binding can be valuable for antiviral drug design), 26 (Pi-stacking [41] and hydrogen bonds [42]) and 39 (Pi-stacking, [39]). To our surprise, according to the SHAP analysis, the formation of hydrogen bonds with residue 22 has a slightly negative effect on the predicted binding activity probability. Earlier literature data indicated that hydrogen bonding with the phosphate oxygen atoms of this residue is important for arginine and argininamide ligand binding [43]. However, the latter molecular dynamics simulations and nuclear magnetic resonance experiments did not support this observation [39], which may explain calculated SHAP values for hydrogen bonds with this residue. The influence of other interactions, such as hydrogen bonding with residues 39 and 24 and lipophilic interactions with residue 27, are harder to interpret due to their mixed contribution to the overall binding activity probability.

SHAP values calculated for the individual ligands may give insights into the contribution of variables (here: interactions) on the probability of the compound being the *binder*, as estimated by the ML model. Selected examples of the binding molecules Mitoxantrone and Amiloride 4h (Fig 4A) show the contributions of the presence or absence of the given interactions to the final prediction of the probability of being a binder for these molecules. Interactions that are colored purple increase the probability of the ligand being the *binder*, while interactions that are colored dark cyan decrease this probability. For Mitoxantrone, the main factors increasing this probability are lipophilic interactions, hydrogen bonds, and cation-anion interactions with residues 40 and 22. Interestingly, this probability is also increased by the fact that the ligand *does not* form hydrogen bonds and lipophilic interaction with residue 27 and hydrogen bonds with residue 24. The probability of the Mitoxantrone being the binder decreases if the ligand does not form the Pi-stacking contact with residue 27. This is in agreement with the literature reports, where the criteria for HIV-1 TAR RNA binders are defined as follows: having an aromatic or heteroaromatic moiety (here, the ligand has this structural feature, but stacking interaction is not observed, thus decreased probability of binding), a cationic residue (the ligand has this structural feature and it has a positive contribution to the overall probability by interactions with residues 22 and 40), and a linker in the structure (a positive contribution to the overall probability by the hydrogen bonding of the hydroxyl group with residue 40) [44,45]. These observations also agree with the interactions detected in a three-dimensional structure of the Mitoxantrone—HIV-1 TAR RNA complex predicted with molecular docking (best-scored pose, Fig 4B). SHAP values calculated for each ligand in the pool and stacked horizontally confirm that, indeed, the probability of binding for the true-inactive ligands is visibly lower than for the real binding ligands (Fig 4C, left and right part of the plot, respectively), although different features are contributing to the final probability values.

**Figure 4.**
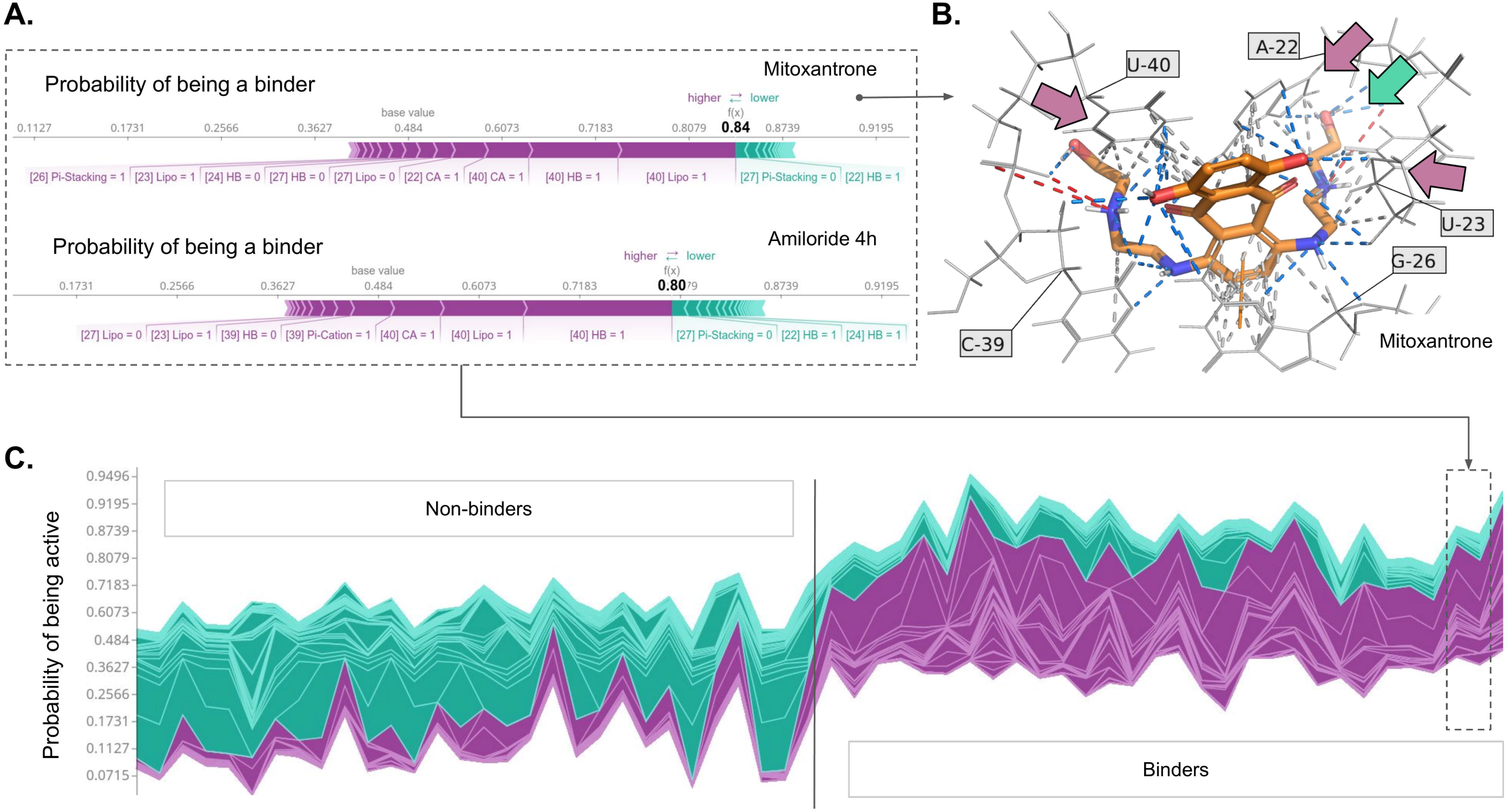
SHAP analysis for the HIV-1 TAR ligand binding activity model trained with the CatBoost classifier on SIFts input. **(A)** Force plots for two selected HIV-1 TAR binding molecules: Mitoxantrone and Amiloride 4h, showing the main features contributing to pushing the model output from the base value (the average model output over the training dataset) to the predicted probability. Features pushing the probability higher are shown in purple, features pushing the probability lower are in dark cyan, and the purple-dark cyan border indicates the probability value. **(B)** Prediction of the binding mode of the Mitoxantrone molecule inside the binding pocket of HIV-1 TAR with detected interactions. The residues forming the interactions with the highest contribution to the model output are indicated by arrows: purple (with the positive contribution) and dark cyan (with the negative contribution). **(C)** SHAP values calculated for each ligand in the pool (with the non-binding ligands located at the first half of the plot and binding ligands at the second half of the plot) stacked horizontally, indicating the predicted probability of the ligand being the *binder*.

Analysis of the variables influencing the decision made by the ML model for Amiloride 4h indicates that the set of interactions is similar to the one for Mitoxantrone, however with some differences (e.g., the importance of the formation of Pi-cation interaction with residue 39, not ranked high in the analysis for Mitoxantrone).

The SHAP method applied to explain ML models built using SIFts can provide valuable insights into the nature of molecular interactions’ importance for small molecule binding. SHAP values of non-covalent interactions combined with analysis of the three-dimensional structure of complexes enable visual interpretation of the ML model and can give instant hints on the nature of the binding phenomena. SHAP analysis not only allowed for the selection of residues and interaction types important for the ligand binding activity but also made it possible to clearly indicate if the given interaction has a positive or a negative influence.

### Summary

Computational methods play a pivotal role in the early stages of drug discovery and are widely applied in virtual screening, structure optimization, and compound activity profiling. With emerging discoveries of the role of RNA in disease, there is a growing interest in utilizing them as potential targets for novel therapies. In this work, we showed that SIFts, combined with ML algorithms, can be readily and effectively used in building predictive models for RNA targets. We showed that these ‘local scoring functions’ (i.e., ML models built with SIFts for a specific molecular target) have superior performance in virtual screening compared to general-purpose scoring functions. On the other hand, to use this approach, a separate ML model must be trained for each molecular target using a dataset of ligands with known activity. Although this requirement currently may be seen as limiting, we believe that with growing interest in studying interactions of small molecules with RNA, the experimental training data will become readily available for many other RNAs, as it is for proteins [46]. Recent applications of protein-focused SIFts-based drug discovery have played an important role in addressing the current needs of global healthcare, such as the COVID-19 pandemic [47,48]. In this work, we also showed that the SIFts-based ML models can be interpreted using modern XAI techniques, such as SHAP, which offer great help in understanding the molecular-recognition process. Merging numerical data with a three-dimensional complex structure can guide medicinal chemists in further directions of developing new potential small molecule drugs. We are convinced that the presented approach will be valuable for the scientific community and facilitate new RNA-based therapies’ discoveries.

## MATERIALS AND METHODS

### Database of RNA-targeting ligands

To construct a dataset containing information on binding and non-binding ligands to the six selected RNA targets, an extensive literature search was performed. Information about the chemical structures of molecules and their binding affinities was collected and tabulated together. As there is a lack of literature reports on non-binding molecules, to simulate the content of the real chemical library where the percentage of binding molecules is low, the additional putative non-binding ligands were generated using the DUD-E methodology (Directory of Useful Decoys: Enhanced; http://dude.docking.org/generate, [24]). The exact procedure was identical to the one described in our previous work [23]. Briefly, ligands whose binding was not described precisely, or was calculated only on the grounds of cell-based assays, were not included in the database. If the RNA sequence, for which the ligand’s binding activity is described, was different from the sequence of experimentally solved structures (Table 1) within the binding site or its proximity up to 8 Å, the ligand was rejected. Structures of the ligands were normalized (i.e., transformed into consistent chemical representation to avoid discrepancies and detect possible errors), and a set of filters was applied to include only drug-like molecules (molecular weight from 90 to 900 daltons, an octanol-water partition coefficient - SlogP from -7 to 9, number of hydrogen bond acceptors up to 18, number of hydrogen bond donors up to 18, number of rotatable bonds up to 18). Next, the PAINS (Pan-assay interference compounds, [49]) filter was applied to exclude ligands for which observed activity was potentially a result of interference with assays and not the interactions with the RNA. Finally, ligands were assigned into one of two groups - *binders* or *non-binders*. The criterion was the binding affinity or biological activity (expressed as K_D_, IC_50_, or K_i_), and ligands with an activity parameter below 300 μM were classified as *binders*. Ligands with an activity parameter higher than 300 μM or described by the authors as “inactive” or “not binding” were classified as *non-binders*. This procedure allowed us to neglect the methodological differences associated with various parameters expressing the activity of compounds and use a binary binding class to compare ligands indirectly. A similar procedure was successfully applied earlier to protein targets (e.g., [50–52]) as well as in our previous work [23]. Afterward, to remove structurally similar compounds, *binders* and *non-binders* were clustered using the *k*-medoids algorithm, and for each cluster, a representative ligand was selected (Fig 5A). Data processing and analysis were performed using the KNIME 4.0.1 analytics platform [53]. The composition of the final datasets is summarized in Table 1.

**Figure 5.**
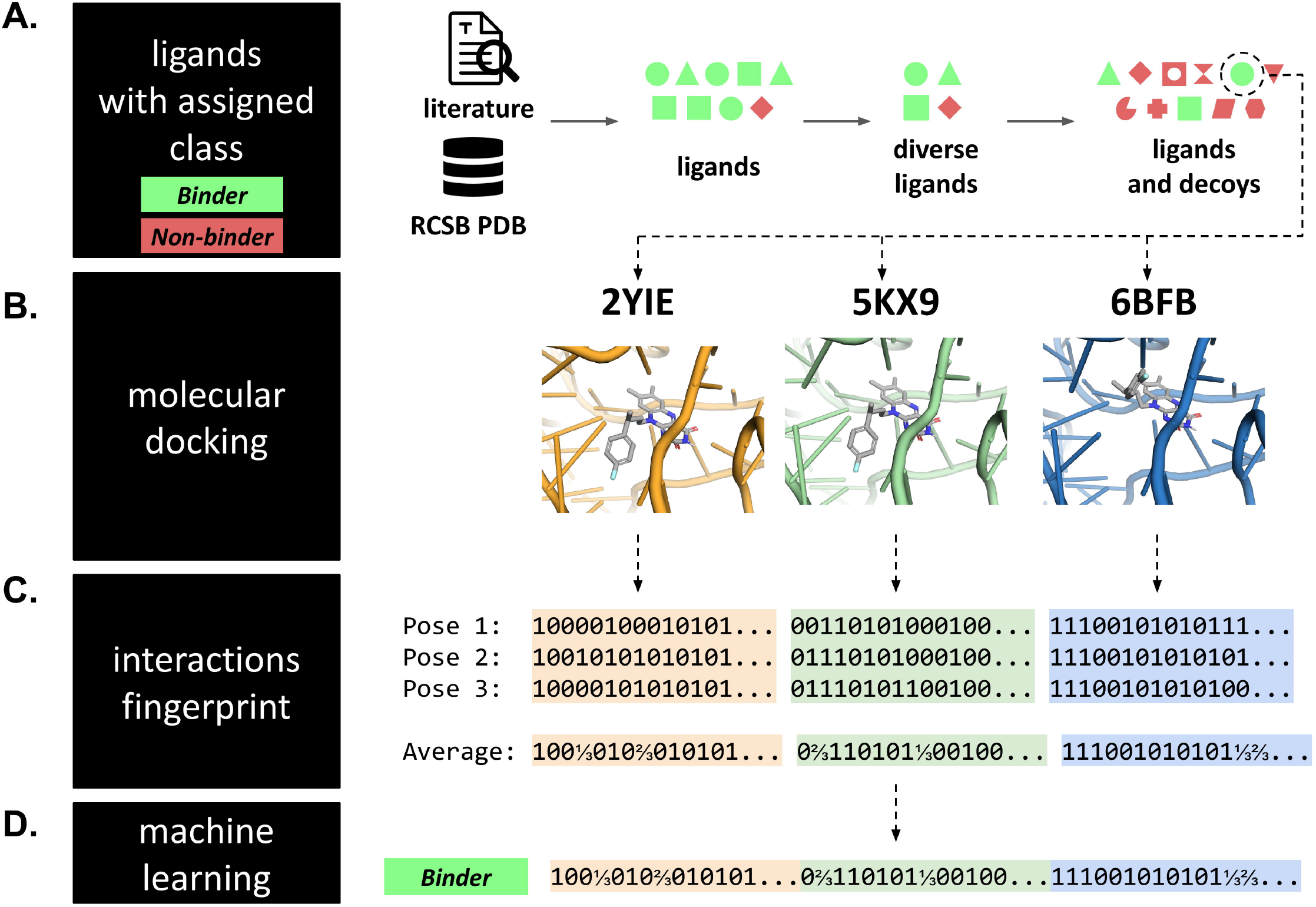
Dataset preparation pipeline. **(A)** For each RNA target, a dataset of ligands with experimentally determined binding to the target RNA was collected from the literature reports and the RCSB PDB database. The pool of ligands was diversified, and putative non-binding ligands were generated. Each molecule from this dataset (with the binary activity class assigned) was **(B)** independently docked to three structures of its corresponding RNA target. **(C)** Next, for each computationally predicted RNA—ligand complex, SIFts were calculated for the three top-scored poses (and then averaged); fingerprints from each complex were concatenated into a single long fingerprint. **(D)** Fingerprints were combined with a binding class of the initial molecule and used as inputs to ML. This procedure was repeated for each molecule in the dataset of a given RNA target, and the resulting fingerprints were collected in a single table, which served as input to ML.

### Datasets for machine learning

The datasets preparation pipeline is summarized in Fig 5.

### Molecular docking

Experimentally solved RNA structures were used for docking small molecule ligands to the six selected viral/bacterial RNA targets (see Table 1, Fig 5B). All macromolecules were preprocessed with the Chimera dockprep pipeline [54]. Docking of small molecule ligands to the aforementioned RNA targets was performed with the rDock docking program (version 2013.1) with dock_solv desolvation potential and a docking radius set to 10 Å [31].

Rescoring of the docked poses was performed using external scoring functions: AnnapuRNA [29], LigandRNA [30], and rf-score-vs [32].

### SIFts’ calculations

Fingerprints type FULL with default settings were calculated for the three top-scored poses of small molecule ligands docked to their RNA target (Fig 5C) using the fingeRNAt software (version 3.0, [23]; fingerprint version FULL; all interactions included) and then averaged. Next, fingerprints calculated for different RNA structures of the same target were horizontally concatenated (this allowed to preserve all information carried by these complexes), and the binding class of the compound was assigned (Fig 5D).

### Machine learning

For building ML models, the mljar-supervised python package was used (https://github.com/mljar/mljar-supervised, version 0.11.1). The tested algorithms included Random Forest, XGBoost, DecisionTree, LightGBM, Extra Trees, and CatBoost. For each method, models were also optimized using feature selection preprocessing (i.e., for the training set, first selecting a subset of important features and next training new models on selected features) and stacking (stacked algorithms are built with predictions from unstacked models). Hyper-parameters were selected using the Optuna optimization framework, with a time budget set to 32400 seconds on 28 CPUs [55]. Additionally, the Baseline predictor, which returns the most frequent label from the training data, was used as a negative control. A 3-fold cross-validation strategy with stratified sampling was used during the optimization, with AUROC as the evaluation metrics.

Feature importance calculation for the CatBoost model built for HIV-1 TAR ligands was performed with CatBoost internal feature importance functions (PredictionValuesChange and LossFunctionChange; CatBoost python module version 1.0.6 [56]). SHAP analysis was performed with the SHAP python module (version 0.41.0, [38,57]); for clarity of data presentation, it was calculated for the top-scored pose with the binding class balanced.

### Data analysis

AUROC values were calculated with the mljar-supervised package (for ML methods) and the scikit-learn python module (version 1.0.1; [58]). The 95% confidence intervals were calculated using a bootstrap algorithm with 1000 iterations using the Seaborn statistical data visualization library (version 0.11.2; [59]). Statistical tests were performed in the SciPy python module (version 1.8.1) using the Wilcoxon signed-rank test, and the two-tailed *p*-values were reported [60].

## DATA AVAILABILITY

The training dataset and the code required for performing all the described analyses can be found at https://github.com/filipsPL/fingernat-ml [25].

## ACKNOWLEDGEMENTS

We thank Dr. Eugene Baulin for his invaluable feedback on the manuscript. This research was carried out in part with the support of the Interdisciplinary Centre for Mathematical and Computational Modelling (ICM) University of Warsaw under computational allocation no G88-1177 to F.S.

This research was funded in part by the National Science Centre, grant number: 2020/39/B/NZ2/03127. For the purpose of Open Access, the author has applied a CC-BY public copyright license to any Author Accepted Manuscript (AAM) version arising from this submission.

